# Host GPCR-cAMP signaling balances Gαs and Gαi activity to control intracellular *Brucella* infection

**DOI:** 10.64898/2026.01.06.697936

**Authors:** Yoon-Suk Kang, James E. Kirby

## Abstract

In this study, we investigated the impact of G protein-coupled receptor (GPCR) signaling on the intracellular replication of the model pathogen *Brucella neotomae*. Building on a prior chemical genetics screen, we identified agonists of the Gαi-coupled adenosine A1 and dopamine D4 receptors as potent inhibitors of intracellular Brucella replication. In contrast, agonists of Gαs-coupled adenosine A2A or dopamine D1 receptors, as well as antagonists of A1 or D4 receptors, either failed to inhibit or enhanced intracellular replication. Wild-type *B. neotomae* induced a rapid, type IV secretion system-dependent increase in host-cell cAMP during early infection. ENBA and cilostamide prevented this infection-associated cAMP increase and completely inhibited intracellular growth; this effect was partially reversed by cell-permeable cAMP analogs Using a real-time NanoBRET biosensor, we detected rapid Gαs activation within minutes of infection that was sustained during wild-type but not ΔvirB4 infection and was abrogated by ENBA or cilostamide. Disruption of early Gαs-cAMP signaling redirected BCVs to replication-incompatible phagolysosomal and autophagy-associated compartments. Collectively, these data support a model in which early GPCR signaling dynamics, balancing Gαs and Gαi pathways, are critical for establishment of productive intracellular Brucella infection.

**Importance Statement:** Brucella species cause chronic infections by surviving and multiplying inside immune cells. To do this, Brucella must remodel the membrane-bound compartment that surrounds it after uptake, steering it away from destructive lysosomes and toward a permissive niche where replication can occur. We found that Brucella rapidly triggers a host signaling response controlled by G protein-coupled receptors, leading to a rise in a common cellular messenger molecule (cAMP) within minutes of infection. This early signal depends on the bacterial type IV secretion system and is required to build the replication-permissive compartment. When we disrupted this signaling with small molecules, bacteria were rerouted into degradative, autophagy-associated compartments and failed to establish productive infection. These results reveal an early host signaling checkpoint that Brucella exploits to create its intracellular niche and suggest that targeting host signaling dynamics, rather than bacterial viability directly, may provide new strategies to block intracellular infection.

## Introduction

*Brucella* species are facultative intracellular bacterial pathogens responsible for brucellosis, a zoonotic disease with substantial human health and agricultural impact worldwide (1). These organisms invade, survive, and replicate within host cells, including macrophages and dendritic cells (2, 3). Intracellular survival enables *Brucella* to evade immune clearance and contributes to chronic bloodstream and reticuloendothelial infection, with severe clinical sequelae such as endocarditis, meningitis, and osteomyelitis. At the cellular level, infection is initiated by bacterial uptake into host cells, followed by trafficking within modified intracellular vacuoles that protect *Brucella* from host antimicrobial defenses (4–6).

Host cells employ multiple defense mechanisms to limit intracellular infection. In addition to pattern recognition receptors (PRRs) such as Toll-like receptors (TLRs) and NOD-like receptors (NLRs), which detect microbial signatures and activate innate immune responses (7), host cells also utilize G protein-coupled receptors (GPCRs) to sense pathogen-associated molecular patterns or infection-induced alterations in cellular physiology (8, 9). GPCR activation engages diverse intracellular signaling pathways, including cAMP production, calcium flux, and MAPK signaling, which can modulate host antimicrobial defenses (10–12).

Despite these defenses, *Brucella* species have evolved strategies to subvert host antimicrobial pathways. These include suppression of autophagy and inflammasome activation and attenuation of reactive oxygen and nitrogen species production within phagolysosomes (4, 13, 14). Many of these effects are mediated through a type IV secretion system (T4SS), which translocates bacterial effector proteins into host cells to remodel intracellular trafficking and signaling pathways.

*Brucella neotomae* shares approximately 99% genetic identity with the major human-pathogenic *Brucella* species (*B. abortus*, *B. suis*, and *B. melitensis*). As reported previously, its intracellular growth properties and murine pathogenic features closely mirror those of human-pathogenic species in cell culture and animal infection models (15). Accordingly, host-pathogen interactions observed using *B. neotomae* are likely to be predictive of those employed by human-pathogenic *Brucella* species. Moreover, as a naturally attenuated pathogen of the desert pack rat, *B. neotomae* provides a lower-risk experimental model that facilitates mechanistic investigation of host-pathogen interactions.

We previously conducted a chemical genetics screen of bioactive compounds that inhibit intracellular replication of *B. neotomae* (6). Notably, several of the most potent inhibitors targeted GPCR-associated signaling pathways, suggesting that GPCR-mediated host signaling may play a critical role during *Brucella* infection. Based on these observations, the aim of the present study was to define how GPCR signaling pathways regulate intracellular *Brucella* replication using the *B. neotomae* infection model.

## Results

### Gαi-coupled GPCR agonists restrict intracellular replication of *B. neotomae*

We previously reported, but did not further explore, a series of G-protein-coupled receptor (GPCR)-related compounds that inhibit intracellular growth of *Brucella neotomae* (Bn) in THP-1 macrophages (6) (**Table S1**). Strong hits were identified as agonists of adenosine A1, dopamine D4, and serotonin 5-HT1A receptors. GPCRs signal through heterotrimeric G proteins, most commonly via Gαs subunits that stimulate adenylate cyclase or Gαi subunits that inhibit adenylate cyclase. Notably, each of the GPCR agonists that inhibited intracellular Bn replication in the initial screen signals through a Gαi-coupled receptor. In contrast, agonists of Gαs-coupled receptors did not inhibit intracellular growth and in some cases potentially stimulated growth. These observations led us to hypothesize that activation of Gαi-coupled GPCR pathways restricts intracellular *Brucella* replication, whereas signaling through Gαs-coupled pathways may be permissive for intracellular growth.

To further analyze the contribution of GPCR signaling to intracellular growth control, *Brucella neotomae* (Bn) containing a chromosomally integrated bacterial *lux* operon was used to infect J774A.1 macrophages. Luminescence output served as a previously validated surrogate marker for intracellular growth based on the robust correlation between luminescence and intracellular colony-forming unit determinations (16). Following infection, cultures were treated with gentamicin to eliminate extracellular bacteria, ensuring that subsequent measurements reflected intracellular replication exclusively (6, 16). Using this experimental system, a panel of GPCR agonists and antagonists was evaluated to define their effects on intracellular growth, quantified by the concentration producing 50% inhibition of intracellular growth (IC_50_) and, where observed, complete inhibition (intracellular MIC).

Agonists of the adenosine A1 receptor (N6-CHA, ENBA, SDZ WAG 994), the dopamine D4 receptor (ABT-724), and the serotonin receptor 5-HT1A, all of which couple to Gαi subunits, potently inhibited intracellular replication, with IC_50_ concentrations in the sub-micromolar range (**Fig. 1A**, **Table 1**). Importantly, concentrations that inhibited intracellular growth were substantially lower than those required to inhibit axenic bacterial growth or host cell viability, supporting a physiological host cell-mediated growth restriction mechanism.

**Figure 1.**
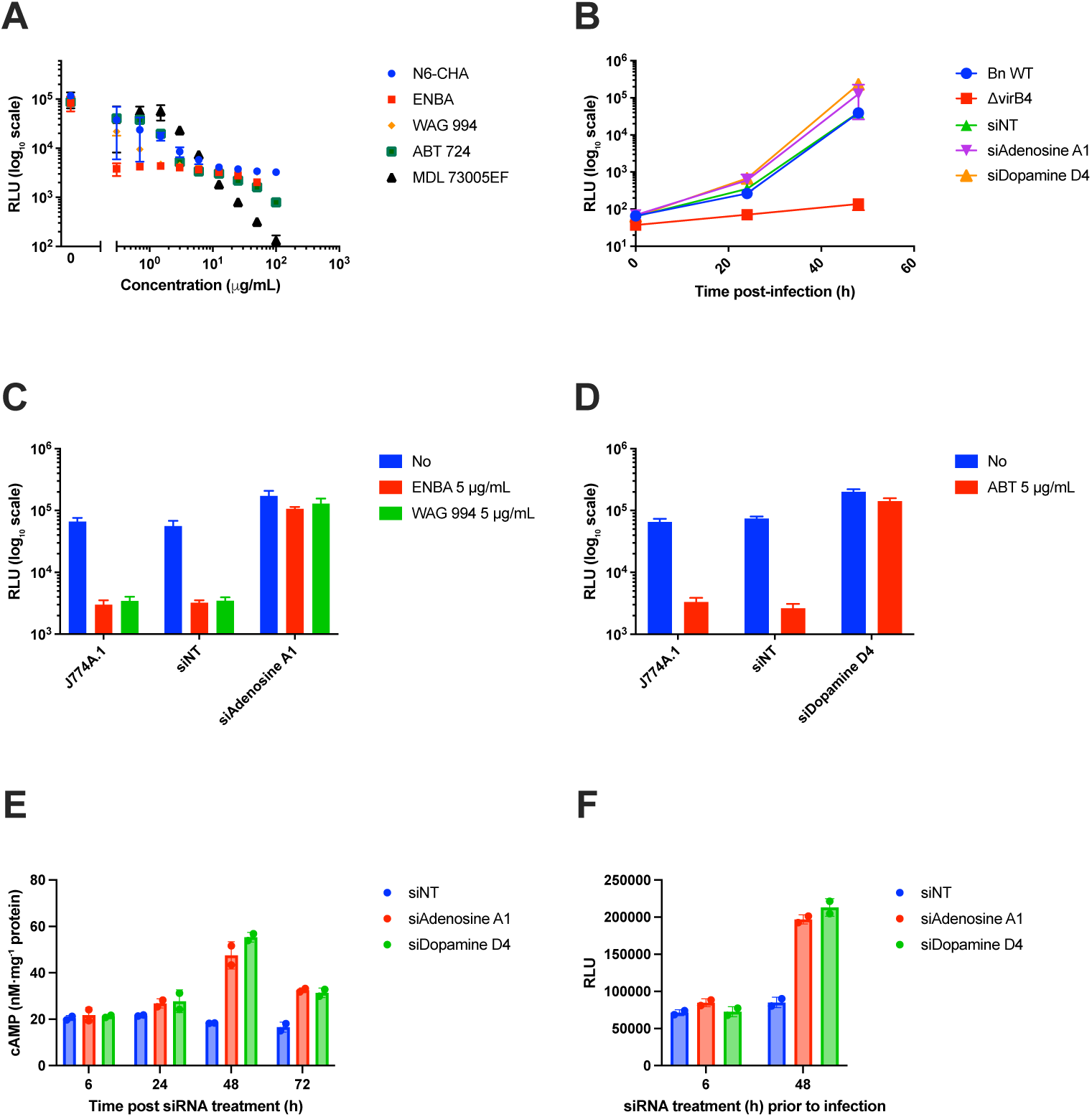
Gαi-coupled host GPCR signaling restricts *Brucella neotomae* intracellular growth via suppression of cAMP. **(A)** Dose-dependent effects of agonists targeting Gαi-coupled host GPCRs on intracellular growth of luminescent *B. neotomae* (Bn) in J774A.1 macrophages measured 48 h post infection. Compounds tested include adenosine A1 receptor agonists N6-cyclohexyladenosine (N6-CHA), ENBA (±-5′-chloro-5′-deoxy-ENBA), and SDZ WAG 994; the dopamine D4 receptor agonist ABT-724; and the serotonin 5-HT1A receptor agonist MDL-73005EF. All receptors targeted in this panel are Gαi-coupled. **(B)** Time course of intracellular growth of wild-type Bn and a type IV secretion-defective mutant (ΔvirB4) in J774A.1 macrophages transfected with non-targeting siRNA (siNT), adenosine A1 receptor siRNA (siA1a), or dopamine D4 receptor siRNA (siD4DR). **(C)** siRNA-mediated knockdown of the adenosine A1 receptor abrogates the intracellular growth-inhibitory effects of the adenosine A1 receptor agonists ENBA and SDZ WAG 994 (each at 5 µg/mL), measured 48 h post infection. **(D)** siRNA-mediated knockdown of the dopamine D4 receptor abolishes the intracellular growth-inhibitory effect of the dopamine D4 receptor agonist ABT-724 (5 µg/mL), measured 48 h post infection. **(E)** Intracellular cAMP concentrations in J774A.1 macrophages following transfection with siNT, siA1a, or siD4DR over a 72-h time course. Knockdown of either Gαi-coupled receptor results in a time-dependent increase in intracellular cAMP. **(F)** Effect of the duration of siRNA pretreatment (6 h vs 48 h) on subsequent intracellular Bn growth measured 48 h post infection. Elevated intracellular cAMP levels at the time of infection (48 h siRNA pretreatment) correlate with increased host cell permissiveness for intracellular bacterial replication. Data shown are the mean ± standard deviation (SD) of at least two independent experiments. Intracellular bacterial burden is reported as relative luminescence units (RLU).

**Table 1.** GPCR agonists that inhibit *Brucella neotomae* intracellular growth in J774A.1 macrophages.

| Compound | Function | IC<sub>50</sub><br>(intracellular) | MIC<sup>a</sup><br>(intracellular) | IC<sub>50</sub><br>(axenic) | MIC<sup>a</sup><br>(axenic) | CC<sub>50</sub><br>J774A.1 | Selectivity<br>(CC<sub>50</sub>/IC<sub>50</sub>)<sup>b</sup> |
| --- | --- | --- | --- | --- | --- | --- | --- |
| N6-Cyclohexyladenosine | A1R agonist (Gαi) | < 0.3 µg/mL | 12.5 µg/mL | > 100 µg/mL | > 100 µg/mL | > 100 µg/mL | >300 |
| (±)-5'-Chloro-5'-deoxy-ENBA | A1R agonist (Gαi) | < 0.3 µg/mL<sup>c</sup> | <0.3 µg/mL | 10.9 µg/mL | > 100 µg/mL | 73 µg/mL | >200 |
| SDZ WAG 994 | A1R agonist (Gαi) | < 0.3 µg/mL | 3 µg/mL | > 100 µg/mL | > 100 µg/mL | > 100 µg/mL | >300 |
| ABT 724 trihydrochloride | D4R agonist (Gαi) | 0.35 µg/mL | 6 µg/mL | 11.7 µg/mL | > 100 µg/mL | > 100 µg/mL | >300 |
| MDL 73005EF hydrochloride | 5-HT1AR agonist (Gαi) | 1.1 µg/mL | 12.5 µg/mL | 11.2 µg/mL | > 100 µg/mL | 75 µg/mL | 75 |

<sup>a</sup>MIC values for intracellular growth represent the lowest concentration tested that resulted in near-complete suppression of intracellular luminescence signal, rather than a CLSI-defined bactericidal endpoint.
<sup>b</sup>All compounds inhibited intracellular replication at concentrations well below host-cell cytotoxicity, consistent with a host-directed mechanism.
<sup>c</sup>IC<sub>50</sub> was below lowest concentration tested.

To further delineate GPCR pathways influencing intracellular replication, additional dopamine and adenosine receptor ligands were examined across a range of concentrations (**Fig. S1**). Among these compounds, only the dopamine D1/D5 receptor (coupled to Gαs) antagonist odapipam inhibited intracellular growth in a dose-dependent manner. In contrast, antagonism of the adenosine A1 receptor with DPCPX resulted in a modest but reproducible increase in intracellular growth, while agonists of the Gαs-coupled adenosine A2A receptors (CGS-21680, regadenoson, UK-432097) and the dopamine D1/D5 receptor agonist SKF38393 had no significant effect on intracellular replication over the concentrations tested. Collectively, these findings indicate that modulation of intracellular growth by GPCR ligands is receptor- and pathway-specific, rather than a nonspecific consequence of GPCR perturbation.

To support the specificity of these pharmacologic observations, the Gαi-coupled adenosine A1 and dopamine D4 receptors were knocked down in host cells prior to infection using small interfering RNA (siRNA) (**Fig. 1B**). J774A.1 macrophages pretreated with non-targeting siRNA (siNT), siA1a (targeting *Adora1*, encoding the adenosine A1 receptor), or siD4 (*Drd4*) *or* siD4DR (targeting *Drd4*) were infected with either wild-type Bn or Bn ΔvirB4, a mutant defective in type IV secretion and unable to establish an intracellular growth niche. At 48 h post-infection, wild-type Bn exhibited significantly enhanced intracellular growth in both siA1a (*P*=0.013) and siD4DR (*P*=0.015) knockdown cells compared with siNT-treated controls. Consistent with receptor-specific effects, the intracellular growth-inhibitory activity of the A1 receptor agonists ENBA and SDZ WAG 994 was completely abrogated by siA1a knockdown (**Fig. 1C**), whereas the inhibitory effect of the D4 receptor agonist ABT-724 was abolished by siD4DR knockdown (**Fig. 1D**). Notably, intracellular growth in siA1a- and siD4DR-treated cells exceeded that observed in siNT-treated controls even in the presence of the corresponding receptor agonists, indicating that tonic signaling through these Gαi-coupled receptors contributes to restriction of intracellular *Brucella* replication.

### Host cell cAMP levels correlate with permissiveness for intracellular *Brucella* growth

Because the adenosine A1 and dopamine D4 receptors are Gαi-coupled GPCRs that, upon activation, inhibit adenylate cyclase and reduce intracellular cAMP levels, we next examined whether intracellular *Bn* growth was influenced by host cell cAMP. We first assessed the time-dependent effects of siRNA-mediated depletion of these receptors on intracellular cAMP concentrations. As shown in **Fig. 1E**, macrophages treated with siA1a or siD4DR exhibited pronounced increases in cAMP, reaching 47.5 and 55.3 nM/mg protein, respectively, at 48 h post-transfection. In contrast, macrophages treated with non-targeting siRNA (siNT) maintained stable cAMP levels of approximately 20 nM/mg protein over the same time course.

To determine whether elevated intracellular cAMP levels at the time of infection were associated with a host cell environment permissive for subsequent intracellular replication, J774A.1 macrophages were infected with *Bn* following siRNA treatment for either 6 h, when cAMP levels remained at baseline, or 48 h, when cAMP levels were elevated. Intracellular bacterial growth was then quantified 48 h post-infection. As shown in **Fig. 1F**, short-term (6 h) treatment with siA1a or siD4DR had no significant effect on intracellular growth compared with the non-targeting siRNA control. In contrast, *Bn* intracellular growth was significantly increased in cells treated with siA1a or siD4DR for 48 h. Together, these results are consistent with a correlation between intracellular cAMP levels at the onset of infection and host cell permissiveness for intracellular bacterial growth.

### Host phosphodiesterase activity regulates intracellular Brucella replication

Gαs-coupled GPCR pathways stimulate adenylate cyclase, leading to increased intracellular cAMP production. Elevated cAMP activates protein kinase A (PKA), which in turn initiates downstream signaling cascades. Phosphodiesterases (PDEs) terminate and modulate cAMP-PKA signaling by hydrolyzing cAMP (and/or cGMP). In a previously reported chemical genetics screen (6), we identified several PDE inhibitors that suppressed *Bn* intracellular replication (**Table S2**).

To further examine the contribution of specific phosphodiesterases, we evaluated the effects of selective and non-selective PDE inhibitors on *B. neotomae* intracellular replication in J774A.1 macrophages (**Fig. 2A**). Both PDE3-selective inhibitors, cilostazol and cilostamide, exhibited strong inhibitory activity, with IC_50_ values of 0.08 µg/mL and 0.02 µg/mL, respectively, and achieved near-complete suppression of intracellular replication at low µg/mL concentrations (**Table 2**). In addition, the PDE7-selective inhibitor BRL50481 produced moderate but incomplete inhibition, with an IC_50_ of 4.1 µg/mL. Similar IC₅₀ values were observed in THP-1 macrophages; however, inhibitory effects plateaued, and complete suppression of intracellular replication was not achieved at the highest concentrations tested. In contrast, neither the non-selective PDE inhibitor IBMX nor inhibitors selective for PDE4 (rolipram) or PDE5 (avanafil) showed substantial inhibition of *B. neotomae* intracellular growth, with IC_50_ values exceeding 50 µg/mL, 21 µg/mL, and 19 µg/mL in J774A.1 cells, respectively.

**Figure 2.**
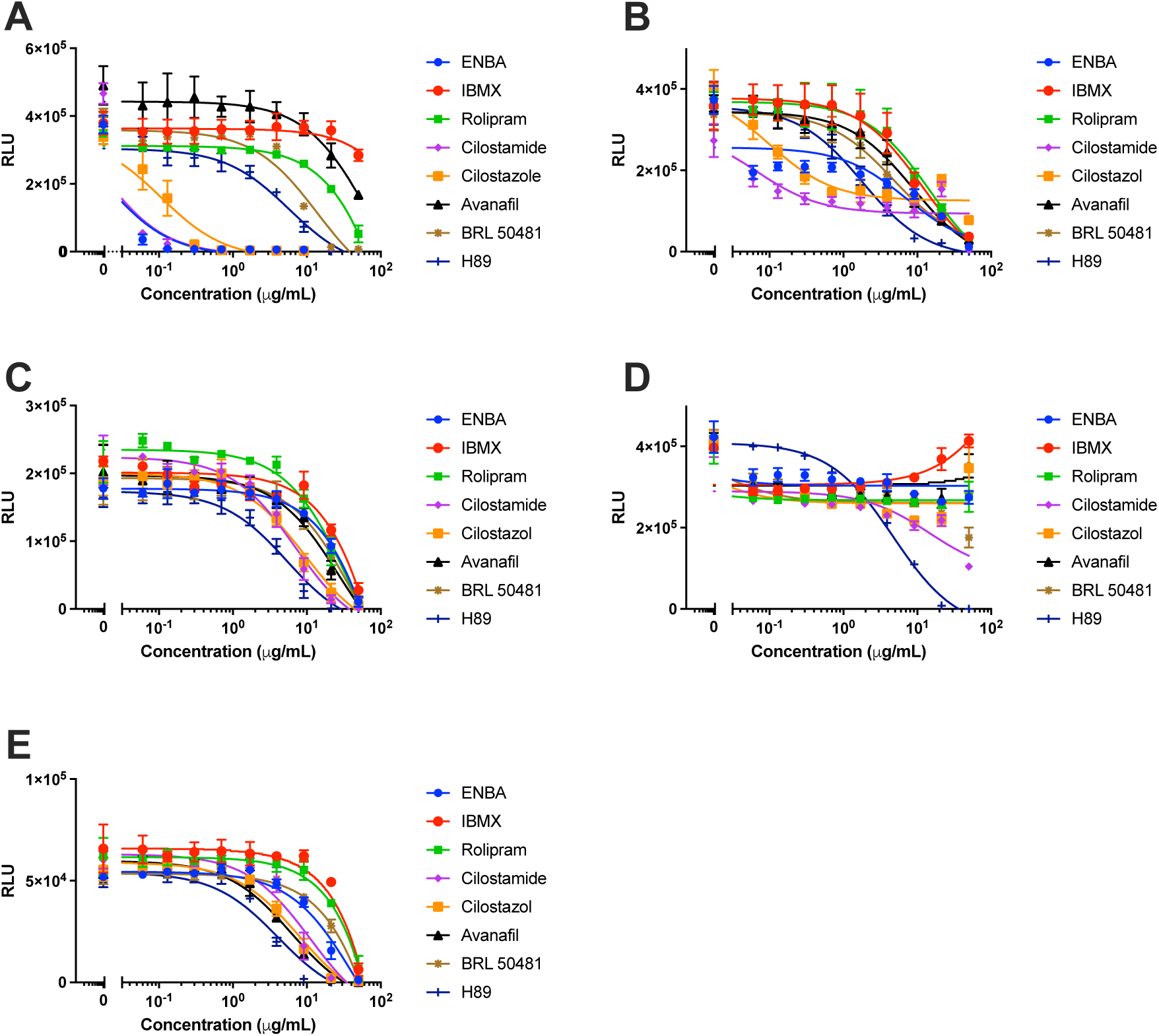
Host phosphodiesterase activity selectively regulates intracellular *Brucella neotomae* replication. **(A)** Intracellular growth of luminescent *B. neotomae* (Bn) in J774A.1 macrophages measured 48 h post infection following treatment with modulators of host cAMP metabolism. Compounds tested include the non-selective phosphodiesterase (PDE) inhibitor IBMX; PDE3-selective inhibitors cilostazol and cilostamide; the PDE4 inhibitor rolipram; the PDE5 inhibitor avanafil; and the PDE7 inhibitor BRL 50481. ENBA, an adenosine A₁ receptor agonist, and H89 are shown for comparison. Data are shown as relative luminescence units (RLU), normalized to untreated controls. **(B)** Effects of the same panel of compounds on intracellular Bn growth in THP-1 macrophages measured 48 h post infection. **(C)** Effects of the indicated compounds on intracellular replication of a lux operon reporter strain of *Legionella pneumophila* in J774A.1 macrophages measured 24 h post infection. Cilostamide was included as a representative PDE3-selective inhibitor. **(D)** Effects of the same compounds on axenic growth of Bn in tryptic soy broth (TSB) medium. **(E)** Effects of the same compounds on axenic growth of *L. pneumophila* in buffered yeast extract (BYE) medium. H89 exhibited comparable inhibitory activity against intracellular and axenic bacteria, consistent with a direct antibacterial effect rather than modulation of host cell signaling. Data shown are the mean ± standard deviation (SD) from two independent experiments.

**Table 2.** Selective inhibition of intracellular *Brucella neotomae* growth by host phosphodiesterase inhibitors.

| IC<sub>50</sub> (μg/mL) | ENBA<sup>a</sup> | IBMX | Rolipram | Cilostamide | Cilostazol | Avanafil | BRL50481 | H89<sup>b</sup> |
| --- | --- | --- | --- | --- | --- | --- | --- | --- |
| Inhibitor specificity | A1R agonist | Non-selective | PDE4 | PDE3 | PDE3 | PDE5 | PDE7 | PKA (non-selective) |
| Bn axenic growth | >50 | >50 | >50 | 15 | >50 | >50 | 20 | 4.7 |
| Intracellular growth, J774A.1 | 0.02 | >50 | 21 | 0.02 | 0.08 | 19 | 4.1 | 5.8 |
| Intracellular growth, THP-1 | 0.35 | 12 | 15 | 0.06 | 0.1 | 8.7 | 4.7 | 1.8 |
| Host cell CC<sub>50</sub> J774A.1 | >50 | >50 | >50 | >50 | >50 | >50 | >50 | >50 |
| Selectivity (CC<sub>50</sub>/IC<sub>50</sub>, intracellular J774A.1)<sup>c</sup> | >2500 | N/A<sup>d</sup> | >2 | >2500 | >600 | >2 | >10 | >9 |

<sup>a</sup>ENBA is included as a comparator that reduces intracellular cAMP through G<sub>ai</sub>-coupled adenosine A<sub>1</sub> receptor activation.
<sup>b</sup>H89 inhibited axenic and intracellular bacterial growth with similar IC<sub>50</sub> values, consistent with a direct antibacterial effect rather than host-directed modulation.
<sup>c</sup> Selectivity was calculated as the ratio of concentrations causing host-cell cytotoxicity (CC<sub>50</sub>) to those inhibiting intracellular *B. neotomae* growth in J774A.1 macrophages. PDE3 inhibitors suppressed intracellular replication at concentrations well below host-cell cytotoxicity, consistent with a host-directed mechanism.
<sup>d</sup>Selectivity was not calculated for IBMX due to lack of measurable intracellular activity at concentrations up to 50 μg/mL. N/A, not applicable.

Importantly, no host-cell cytotoxicity was observed for any compound at concentrations up to 50 µg/mL (data not shown), and active inhibitors were highly selective for intracellular replication relative to axenic bacterial growth (**Fig. 2A, D**; **Table 2**). In contrast, none of the PDE inhibitors selectively inhibited intracellular replication of the type IV secretion system-dependent pathogen *L. pneumophila* in either J774A.1 or THP-1 macrophages, as reflected by comparable IC_50_ values for inhibition of axenic and intracellular growth (**Table 3**; **Fig. 2C, E**). Inhibition observed at the highest concentrations tested was therefore attributable to direct antibacterial effects rather than host-directed modulation.

**Table 3.** Lack of selective inhibition of *Legionella pneumophila* growth by host phosphodiesterase inhibitors.

| IC<sub>50</sub> (µg/mL) | ENBA<sup>a</sup> | IBMX | Rolipram | Cilostamide | Cilostazol | Avanafil | BRL50481 | H89<sup>b</sup> |
| --- | --- | --- | --- | --- | --- | --- | --- | --- |
| Inhibitor specificity | A1R agonist | Non-selective | PDE4 | PDE3 | PDE3 | PDE5 | PDE7 | PKA |
| Lp axenic growth | 16 | 35 | 33 | 7.4 | 6.9 | 6.4 | 22 | 4.1 |
| Intracellular growth, J774A.1 | 21 | 22 | 20 | 6.0 | 9.7 | 17 | 17 | 5.1 |

<sup>a</sup>ENBA is included as a comparator that reduces intracellular cAMP through G<sub>ai</sub>-coupled adenosine A1 receptor activation.
<sup>b</sup>H89 inhibited axenic and intracellular bacterial growth with similar IC<sub>50</sub> values, consistent with a direct antibacterial effect rather than host-directed modulation.

Taken together, these findings identify host cell PDE3, and to a lesser extent PDE7, as specific regulators of the *Brucella* intracellular growth niche, while highlighting the pathogen-specific nature of host-directed PDE modulation.

### cAMP signaling promotes early intracellular replication of *B. neotomae*

One puzzling observation was that among the most potent inhibitors of intracellular *B. neotomae* replication were the Gαi-coupled GPCR agonist ENBA and the PDE3 inhibitors cilostazol and cilostamide, despite these reagents being generally thought to regulate intracellular cAMP levels in opposing directions (17, 18). To investigate this apparent paradox, we infected J774A.1 macrophages with Bn and measured intracellular cAMP concentrations in the presence of compounds that either inhibited (ENBA, cilostamide) or failed to inhibit (IBMX, rolipram, avanafil) intracellular growth (**Fig. 3A**). Infection with wild-type Bn, but not the ΔvirB4 mutant, resulted in significantly increased intracellular cAMP levels, indicating that cAMP induction is dependent on a functional type IV secretion system. Notably, both ENBA and cilostamide abrogated this infection-associated cAMP increase, whereas PDE inhibitors that did not inhibit intracellular growth had no significant effect.

**Figure 3.**
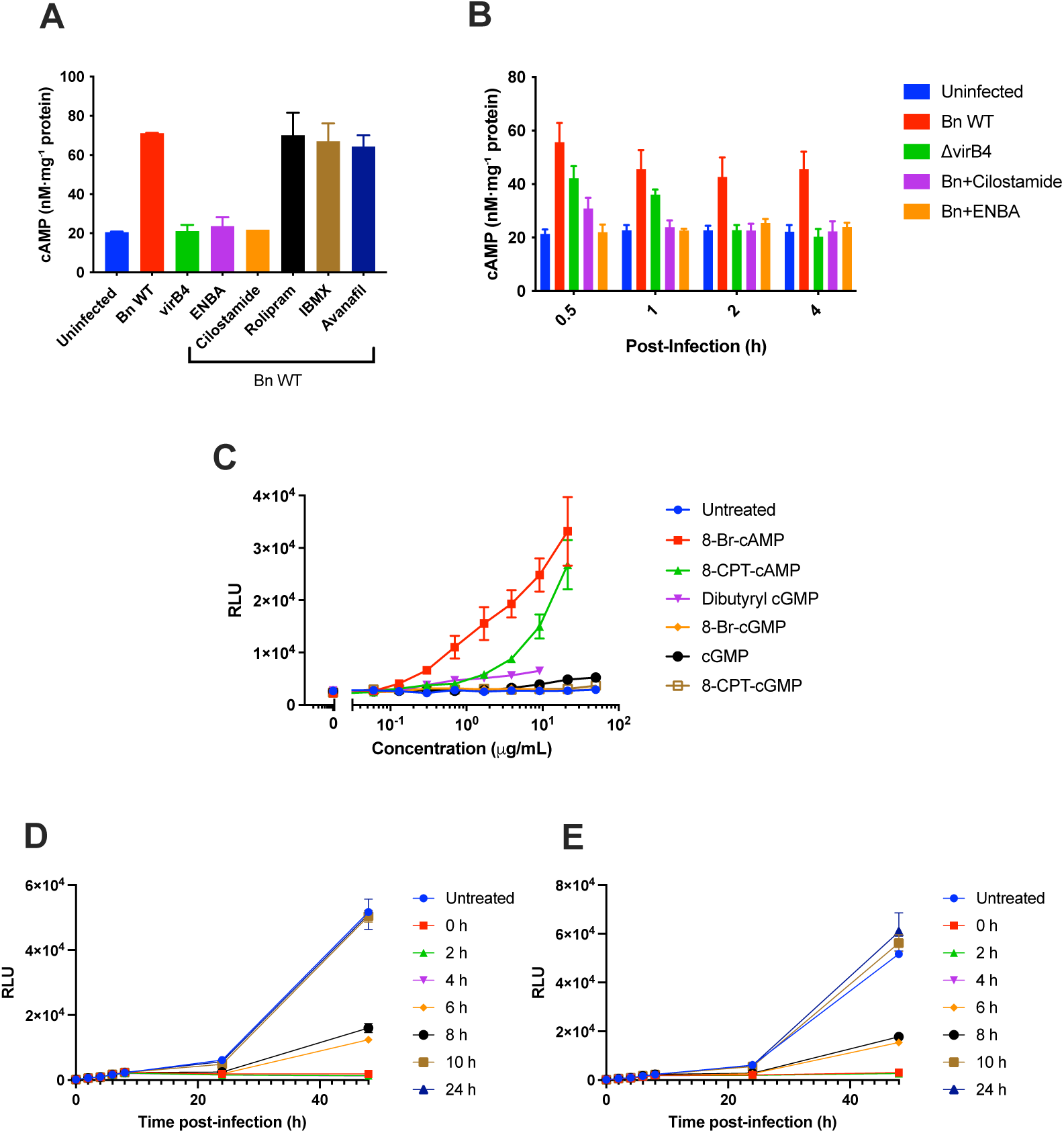
Early activation of host cAMP signaling promotes intracellular replication of *Brucella neotomae*. **(A)** Intracellular cAMP concentrations in J774A.1 macrophages measured 48 h post infection with wild-type *B. neotomae* (Bn) in the presence of the indicated phosphodiesterase (PDE) inhibitors or the adenosine A1 receptor agonist ENBA (each at 2 µg/mL). cAMP levels were normalized to total protein. Uninfected and ΔvirB4-infected cells are shown for comparison. **(B)** Time course of intracellular cAMP levels during early infection with wild-type Bn or the type IV secretion system-defective ΔvirB4 mutant. Where indicated, cells were treated with ENBA or the PDE3 inhibitor cilostamide (2 µg/mL). Data represent mean ± SD from three independent experiments. **(C)** Rescue of ENBA-mediated inhibition of intracellular Bn growth by cell-permeable cyclic nucleotide analogs. J774A.1 macrophages were infected in the presence of ENBA (2 µg/mL) and treated with the indicated cAMP or cGMP analogs at concentrations below axenic growth IC_50_ and host cell cytotoxicity thresholds (**Table S3**). Intracellular growth was quantified 48 h post infection and expressed as relative luminescence units (RLU). **(D)** Effect of delayed addition of ENBA (2 µg/mL) on intracellular Bn growth. ENBA was added at the indicated times post infection, and intracellular growth was monitored over 48 h using a luminescence reporter. **(E)** Effect of delayed addition of cilostamide (2 µg/mL) on intracellular Bn growth, performed as in panel D. In panels D and E, the increase in luminescence observed during the first 6-8 h post infection reflects early metabolic adaptation associated with establishment of the intracellular niche rather than bacterial replication, as intracellular CFU counts do not increase during this period (data not shown). Data shown are the mean ± SD of three independent experiments.

To further define the temporal dynamics of cAMP signaling during infection, we examined intracellular cAMP levels during the early stages following bacterial uptake (**Fig. 3B**). Intracellular cAMP concentrations peaked within 30 min of infection and subsequently declined modestly but remained above baseline. Infection with the ΔvirB4 mutant or treatment with cilostamide resulted in a blunted cAMP increase at 30 min that returned to baseline by 1-2 h post infection, whereas ENBA blocked the cAMP increase at the earliest time point examined. These results suggest that wild-type Bn actively stimulates host cAMP signaling during the initial stages of infection.

The reduction in intracellular cAMP observed in cilostamide-treated, infected cells was unexpected given the established role of PDE3 inhibitors in blocking cAMP degradation.

However, in uninfected J774A.1 macrophages, cilostamide treatment alone increased intracellular cAMP approximately twofold within 30 min, and this elevation was maintained for at least 2 h (**Fig. S2**). Thus, the effect of cilostamide on cAMP during infection appears to reflect interference with infection-induced cAMP signaling rather than suppression of basal host cAMP levels.

These findings led us to ask whether inhibition of intracellular replication by ENBA could be rescued by restoring early cAMP signaling. To test this, ENBA-treated macrophages were supplemented with membrane-permeable cAMP or cGMP analogs (Fig. 3C). cAMP analogs (8-Br-cAMP and 8-CPT-cAMP) robustly restored intracellular growth, producing up to ∼15-fold and ∼10-fold increases, respectively, at concentrations that did not affect axenic bacterial growth or host cell viability (**Table S3**). In contrast, cGMP analogs produced minimal or no rescue. These results indicate that early host cAMP signaling directly contributes to establishment of a permissive intracellular niche.

Protein kinase A (PKA) has previously been implicated in *Brucella* intracellular replication based on studies using the inhibitor H89 (19). However, in our system, H89 inhibited intracellular and axenic Bn growth to a similar extent in both J774A.1 and THP-1 macrophages, indicating a direct antibacterial effect rather than host-directed signaling. Moreover, treatment with the highly specific PKA inhibitor, myristolated PKI 14-22 amide, did not affect intracellular Bn growth (**Fig. S3**). Together, these observations indicate that PKA activity is not required for intracellular Bn replication and that cAMP influences infection through PKA-independent pathways.

Finally, to define when during infection GPCR-cAMP signaling is required, we added ENBA or cilostamide at defined times post infection (**Fig. 3D, E**). Addition of either compound within the first 4-6 h post infection markedly inhibited intracellular growth, whereas delayed addition at 10 h or later had little effect. Because luminescence increases during the first 6-8 h reflect metabolic adaptation rather than bacterial replication, these results suggest that ENBA and cilostamide disrupt host cell processes required for establishment of the intracellular niche during the early stages of infection, prior to formation of the ER-derived replicative vacuole.

### Sustained Gαs activation during *Brucella neotomae* infection requires type IV secretion

Based on the growth-inhibitory effects of Gαi-coupled GPCR agonists and the growth-promoting effects of selected Gαs agonists, we next asked whether *Brucella neotomae* (Bn) actively promotes host Gαs signaling to support intracellular replication. In its inactive state, heterotrimeric G proteins consist of a Gα subunit associated with Gβγ and coupled to a GPCR. Upon receptor activation, GDP-GTP exchange on Gαs leads to dissociation of Gαs from Gβγ, enabling downstream signaling through effectors such as adenylyl cyclase and MAP kinase pathways. We previously demonstrated MAP kinase activation downstream of GPCR signaling during Bn infection (6).

To directly monitor Gαs activation in live cells, we employed a NanoBRET (bioluminescence resonance energy transfer)-based assay in which Gαs is fused to NanoLuc luciferase and Gγ to a YFP reporter, analogous to previously described systems for monitoring GPCR-mediated G protein activation (20, 21), with minor modifications (**Fig. 4A**). In this system, the inactive heterotrimeric complex yields a high BRET signal due to close proximity of Gαs-NanoLuc and Gγ-YFP, whereas Gαs activation and dissociation from Gβγ result in a decrease in BRET signal.

**Figure 4.**
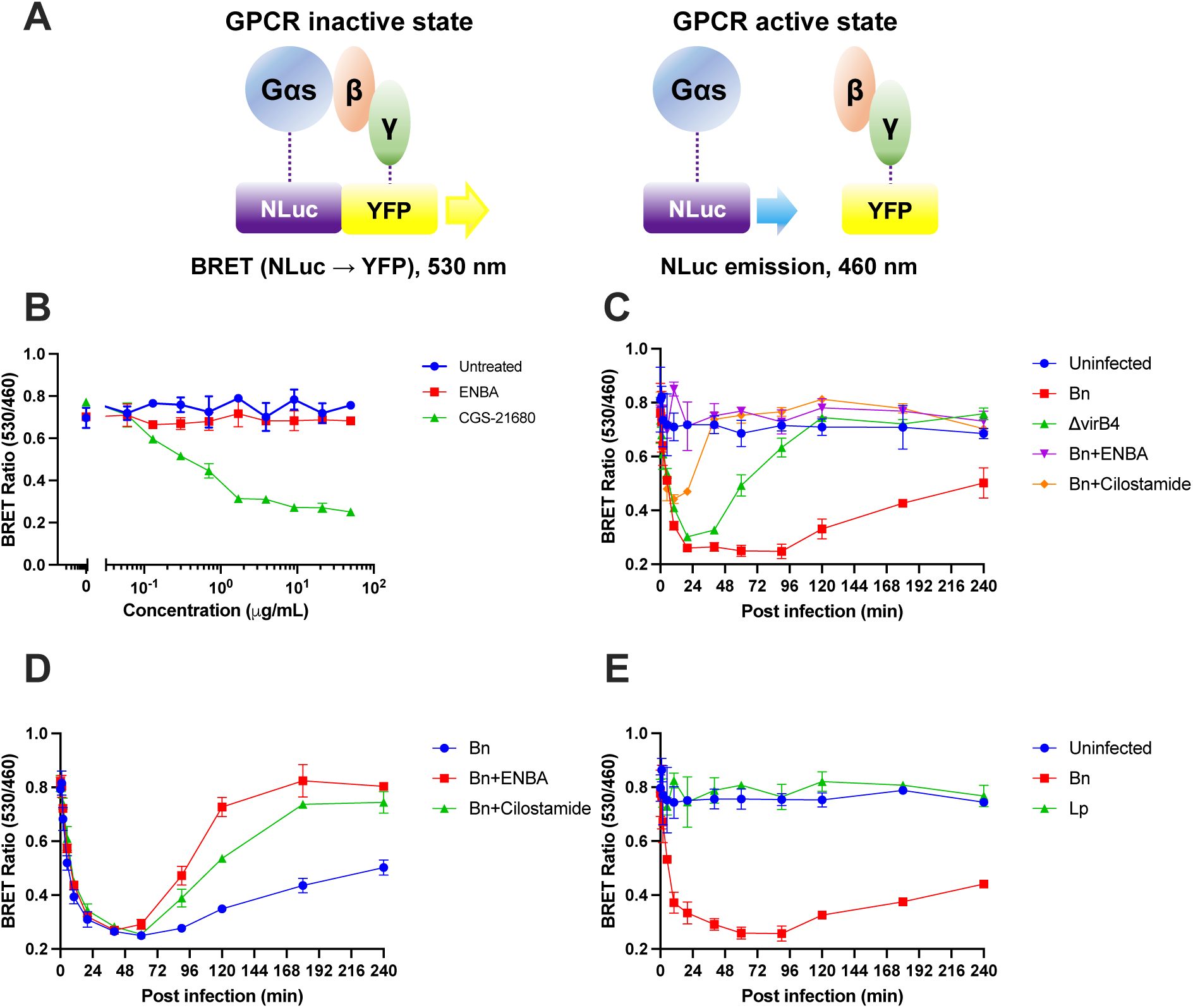
Sustained Gαs activation during *Brucella neotomae* infection requires type IV secretion. **(A**) Schematic of the NanoBRET assay used to monitor Gαs activation in live cells. In the inactive state, Gαs fused to NanoLuc luciferase (NLuc) remains associated with Gβγ, with Gγ fused to YFP, resulting in a high BRET signal due to energy transfer from NLuc emission (460 nm) to YFP emission (530 nm). Upon Gαs activation and dissociation from Gβγ, BRET efficiency decreases, and donor emission at 460 nm predominates. **(B)** Validation of the NanoBRET assay using GPCR agonists with defined G-protein coupling. Treatment with the adenosine A₂A receptor agonist CGS-21680 (Gαs-coupled), but not the adenosine A₁ receptor agonist ENBA (Gαi-coupled), resulted in a concentration-dependent decrease in BRET signal measured 30 min after agonist addition, consistent with selective activation of Gαs. **(C)** Real-time monitoring of Gαs activation during infection of J774A.1 macrophages with wild-type *B. neotomae* (Bn) or the type IV secretion-deficient ΔvirB4 mutant. Where indicated, ENBA or the PDE3 inhibitor cilostamide (each at 2 µg/mL) was added at the time of infection. Uninfected cells are shown as a negative control. Sustained Gαs activation was observed during wild-type Bn infection but not during ΔvirB4 infection and was blocked by ENBA or cilostamide. **(D)** Effect of delayed addition of ENBA or cilostamide (2 µg/mL) at 1 h post infection on Gαs activation during wild-type Bn infection. Addition of either compound resulted in rapid inactivation of Gαs and return of the BRET signal toward baseline. **(E)** Gαs activation during infection with *Legionella pneumophila* (Lp). In contrast to Bn, Lp infection did not induce detectable Gαs activation. For all panels, BRET values are expressed as the ratio of emission at 530 nm to 460 nm. Data points represent the mean ± SD of two independent biological experiments.

We first validated the specificity of this NanoBRET assay using GPCR agonists with defined G-protein coupling. Treatment of transfected host cells with the adenosine A2A receptor agonist CGS-21680, which signals through Gαs, produced a concentration-dependent decrease in BRET signal measured 30 min after agonist addition (**Fig. 4B**). In contrast, treatment with the adenosine A1 receptor agonist ENBA, which signals through Gαi, had no effect on the BRET signal, confirming selective detection of Gαs activation.

We next applied this assay to examine Gαs activation during infection. Infection of J774A.1 macrophages with wild-type Bn resulted in rapid Gαs activation within minutes of bacterial uptake (**Fig. 4C**). Infection with the type IV secretion-deficient ΔvirB4 mutant also triggered an initial decrease in BRET signal, indicating early Gαs activation; however, this response was transient and rapidly returned toward baseline. In contrast, wild-type Bn infection produced sustained Gαs activation over the duration of the assay. This early, transient activation may reflect host cell responses to bacterial uptake, whereas sustained Gαs activation requires active type IV secretion-dependent modulation. These results indicate that while initial Gαs activation may occur independently of type IV secretion, sustained activation requires an intact type IV secretion system.

We next assessed whether pharmacologic modulation of GPCR signaling affected Gαs activation during infection. Addition of the Gαi-coupled GPCR agonist ENBA at the time of infection completely blocked Gαs activation during wild-type Bn infection (**Fig. 4C**). Treatment with the PDE3 inhibitor cilostamide permitted an initial activation of Gαs but was followed by a rapid return of the BRET signal toward baseline, consistent with loss of sustained activation. These NanoBRET results closely parallel the effects of ENBA and cilostamide on intracellular cAMP dynamics observed during early infection (**Fig. 3B**).

To further probe regulation of Gαs signaling during established infection, ENBA or cilostamide was added to wild-type Bn-infected macrophages at 1 h post infection (**Fig. 4D**). Addition of either compound resulted in rapid inactivation of Gαs and restoration of the BRET signal toward baseline levels. ENBA-mediated activation of Gαi may therefore antagonize Gαs signaling, potentially by promoting reassociation of Gαs with Gβγ or accelerating GTP hydrolysis, whereas cilostamide-induced inactivation of Gαs may occur through mechanisms independent of direct Gαi activation.

Finally, to assess whether Gαs activation represents a general feature of intracellular bacterial infection, we examined Gαs activation during infection with *Legionella pneumophila* (Lp02fla). In contrast to Bn, Lp infection did not induce detectable Gαs activation (**Fig. 4E**). Consistent with this observation, intracellular replication of *L. pneumophila* was unaffected by ENBA or cilostamide treatment. Together, these results demonstrate that sustained Gαs activation during infection is specific to *Brucella neotomae*, requires a functional type IV secretion system, and correlates with host cAMP signaling and intracellular replication, supporting a model in which Bn actively promotes host Gαs signaling to establish a permissive intracellular replication niche.

### cAMP signaling is required for maturation of the replicative Brucella-containing vacuole

Following uptake by host macrophages, *Brucella* occupies a series of temporally and spatially distinct intracellular compartments, transitioning from an endocytic vacuole to an ER-derived replicative niche beginning approximately 6-12 h post infection (3, 16, 22). Perturbation of GPCR-cAMP signaling early during infection strongly inhibits intracellular replication, prompting us to ask whether this growth restriction reflects impaired maturation of the Brucella-containing vacuole (BCV).

To assess early trafficking events, we quantified BCV association with Rab7, a marker of late endosomes and lysosomes. During the first 1-4 h post infection, BCVs exhibited similarly high Rab7 association (approximately 50-70%) under all conditions examined (Fig. 5A-C). However, by 12 h and at later time points, BCV trafficking diverged. In untreated wild-type infection, Rab7 association progressively declined, consistent with exit from the endolysosomal pathway during establishment of the replicative niche. In contrast, Rab7 association remained persistently elevated in ENBA- and cilostamide-treated cells, indicating a failure to progress beyond late endosomal compartments.

**Figure 5.**
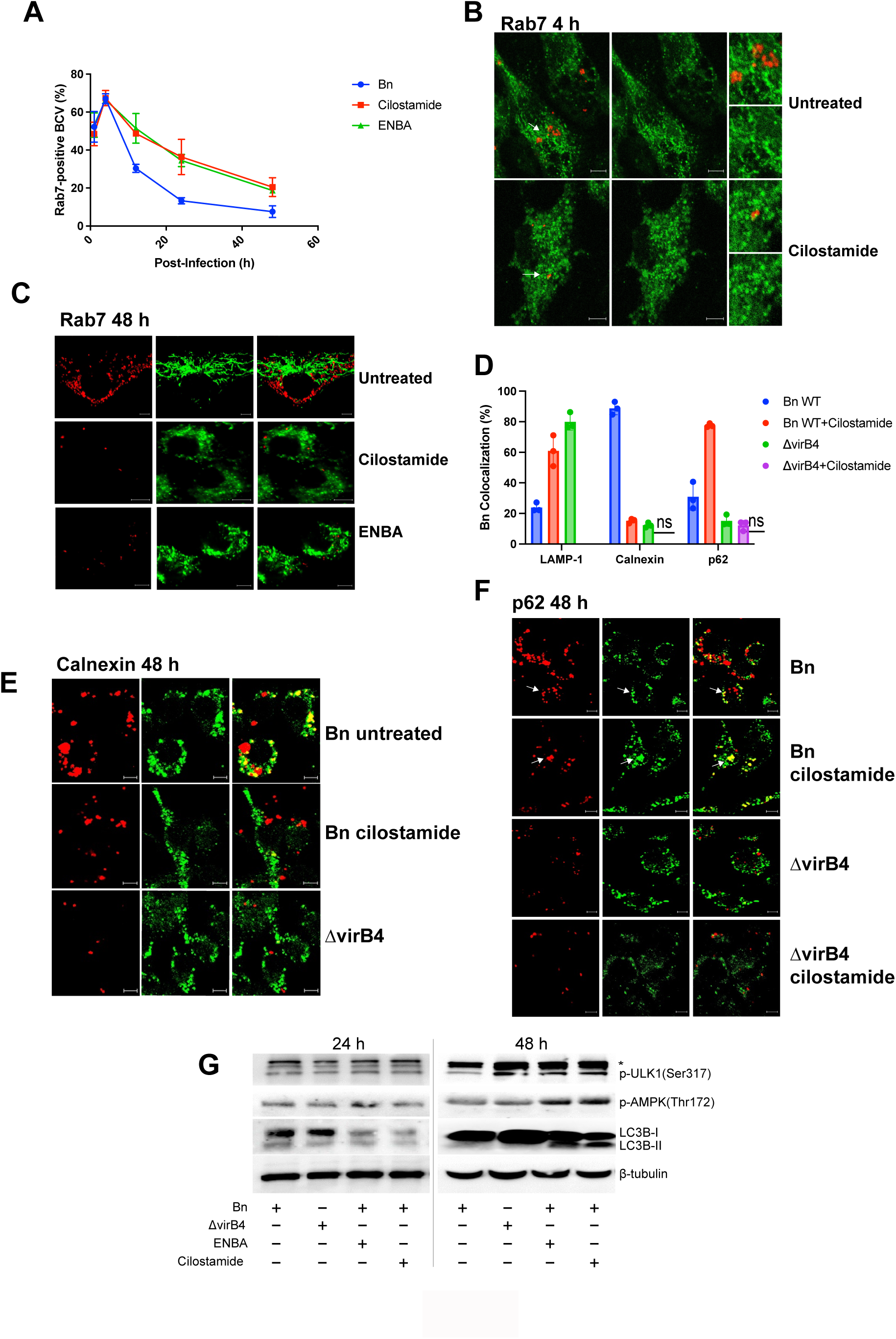
Disruption of host cAMP signaling impairs maturation of the replicative Brucella-containing vacuole. **(A)** Quantification of Rab7 association with *Brucella neotomae*-containing vacuoles (BCVs) over time. J774A.1 macrophages were infected with tdTomato-expressing wild-type Bn in the presence or absence of ENBA or cilostamide (2 µg/mL), and BCV colocalization with the late endosomal marker Rab7 was assessed by confocal microscopy at the indicated time points. Data represent mean ± SD from at least 50 BCVs per condition from three independent experiments. **(B, C)** Representative confocal images showing Rab7 colocalization with BCVs during infection with wild-type Bn at 4 h (B) and 48 h (C) post infection, under the indicated treatment conditions. Bn (red) were labeled with a tdTomato fluorescent reporter for visualization, and Rab7 was identified by immunofluorescence (green). White arrows indicate Rab7-positive BCVs enlarged in insets. Scale bars, 5 µm. **(D)** Quantification of colocalization of BCVs with LAMP-1 (lysosomal marker), calnexin (endoplasmic reticulum marker of replicative BCVs), and p62 (autophagy adaptor protein) at 48 h post infection. J774A.1 cells were infected with tdTomato-expressing wild-type Bn or the ΔvirB4 mutant in the presence or absence of cilostamide (2 µg/mL). Data represent mean ± SD from at least 50 BCVs per condition from three independent experiments. Statistical significance was determined by two-way ANOVA with Dunnett’s multiple-comparison test. **(E, F)** Representative confocal images showing BCV (red) colocalization with calnexin (E) and p62 (F) (green) at 48 h post infection. Untreated wild-type Bn localizes predominantly to calnexin-positive ER-derived replicative BCVs, whereas cilostamide treatment increases association with p62-positive compartments (white arrows indicate a representative example of colocalization, appearing yellow in merged images). ΔvirB4 BCVs show high lysosomal association and minimal calnexin or p62 colocalization. Scale bars, 5 µm. **(G)** Immunoblot analysis of autophagy-associated signaling pathways at 24 h and 48 h post-infection. J774A.1 macrophages infected with wild-type *Brucella neotomae* (Bn) or the ΔvirB4 mutant, in the presence or absence of ENBA or cilostamide (2 µg/mL), were analyzed for phosphorylation of AMPK (Thr172) and ULK1 (Ser317), as well as LC3B processing. Phosphorylation of AMPK and ULK1 was most apparent at 48 h post-infection and was enhanced in inhibitor-treated cells. Conversion of LC3B-I to LC3B-II was minimal during infection with wild-type Bn or the ΔvirB4 mutant alone, but inhibitor treatment during wild-type infection was associated with reduced LC3B-I abundance and increased relative LC3B-II accumulation, most evident at 48 h. β-tubulin served as a loading control; asterisks indicate non-specific bands.

We next examined BCV identity at later stages of infection using markers associated with lysosomes, autophagy-related compartments, and ER-derived replicative vacuoles. At 48 h post-infection, untreated wild-type BCVs showed low colocalization with LAMP-1, high colocalization with the ER marker calnexin, and minimal association with p62, consistent with maturation into ER-derived replicative BCVs (**Fig. 5D-F**). As expected, ΔvirB4 BCVs displayed increased LAMP-1 and reduced calnexin colocalization, consistent with diversion to a degradative lysosomal pathway. Notably, cilostamide treatment shifted wild-type BCVs toward a distinct phenotype characterized by increased association with both LAMP-1 and p62 and reduced colocalization with calnexin, consistent with trafficking into autophagolysosomal compartments. Under these conditions, intracellular replication was severely restricted, with BCVs containing only single bacteria or small aggregates.

To determine whether these alterations in BCV trafficking were accompanied by changes in autophagy-associated signaling pathways, we examined phosphorylation and processing of key regulators of autophagy. At 24 h and more prominently at 48 h post-infection, *B. neotomae*-infected cells treated with ENBA or cilostamide exhibited increased phosphorylation of AMPK (Thr172) and ULK1 (Ser317) relative to untreated wild-type infection (**Fig. 5G**). In contrast, infection with wild-type *B. neotomae* or the type IV secretion system-deficient ΔvirB4 mutant alone was associated with minimal LC3B processing. Notably, ENBA or cilostamide treatment during wild-type infection was associated with reduced abundance of LC3B-I and increased relative accumulation of LC3B-II, most evident at 48 h, whereas LC3B-II was not detected under ΔvirB4 infection conditions, with or without inhibitor treatment (**Fig. 5G; Fig. S4**). Together, these findings indicate that pharmacologic disruption of cAMP signaling during wild-type infection promotes engagement of autophagy-associated signaling and LC3B processing in parallel with altered BCV trafficking and failure to establish an ER-derived replicative niche.

Consistent with our prior work, sustained phosphorylation of p38 MAP kinase was observed during wild-type but not ΔvirB4 infection at 48 h post infection (**Fig. S5**). Notably, p38 phosphorylation during wild-type infection was reduced by ENBA or cilostamide treatment at late time points, placing p38 activation downstream of inhibitor-sensitive signaling events and providing temporal context for the GPCR-cAMP signaling cascade described here and previously (6). Together, these findings support a model in which early cAMP signaling promotes maturation of the ER-associated replicative BCV, whereas pharmacologic disruption of this pathway diverts wild-type *Brucella* into LAMP-1- and p62-positive compartments with autophagic features, resulting in failure to establish a productive intracellular replicative niche.

## Discussion

The ability of *Brucella* species to survive and replicate within host cells, particularly macrophages, is central to their pathogenesis. Intracellular replication depends on successful maturation of the Brucella-containing vacuole (BCV) into an endoplasmic reticulum (ER)- derived replicative niche, a process that requires a functional type IV secretion system (T4SS). Mutants defective in T4SS-mediated intracellular replication are avirulent and rapidly cleared in experimental animal models, underscoring the importance of host-pathogen interactions that govern BCV maturation and intracellular survival (23, 24). Despite substantial progress in identifying T4SS effector proteins that contribute to these processes, the host signaling pathways exploited by *Brucella* to establish the intracellular replicative niche have remained incompletely understood.

In prior work, we reported a chemical genetics screen that identified host-directed small molecules capable of inhibiting intracellular replication of *Brucella neotomae* (Bn) in macrophages (15). Among the compounds identified were inhibitors targeting host signaling pathways, including MAP kinase signaling, which disrupted BCV maturation and intracellular growth. In that study, Bn infection was associated with robust and sustained activation of p38 MAP kinase alongside suppression of ERK and MEK phosphorylation, highlighting the importance of host signaling modulation during establishment of the intracellular niche. However, the upstream host pathways responsible for initiating these signaling events during infection were not defined.

A notable feature of the chemical genetics screen was the enrichment of compounds targeting G protein-coupled receptors (GPCRs), including agonists of GPCRs associated with Gαi subunits and antagonists of GPCRs associated with Gαs subunits. This pattern suggested that the balance of GPCR-mediated signaling influences the ability of *Brucella* to replicate within macrophages and raised the possibility that the pathogen exploits GPCR pathways during early stages of infection. GPCR signaling can function upstream of diverse intracellular signaling cascades in macrophages, including cyclic AMP (cAMP)-dependent pathways downstream of Gαs and signaling mediated by Gβγ subunits, providing a plausible link between GPCR engagement and the host responses observed during *Brucella* infection.

GPCRs are central regulators of immune cell function, mediating responses to extracellular metabolites, neurotransmitters, and inflammatory mediators, and play key roles in inflammation and innate immune defense (9, 25, 26). Several bacterial pathogens exploit GPCR pathways to modulate host immune responses and promote intracellular survival. For example, *Staphylococcus aureus* activates the anti-inflammatory adenosine A2A receptor to suppress neutrophil activation, while bacterial leukotoxins directly target chemokine receptors to interfere with host signaling (27, 28). In the context of *Brucella*, *B. abortus* has been reported to induce transcriptional upregulation of the adenosine A2B receptor during macrophage infection; however, pharmacologic modulation of this receptor failed to alter intracellular bacterial burden at later time points (29, 30), highlighting the importance of temporal and contextual aspects of GPCR signaling during infection.

Adenosine and dopamine receptors are established regulators of cAMP signaling, controlling intracellular cAMP levels and downstream phosphorylation cascades (31–33). In the present study, pharmacologic activation of Gαs-coupled GPCRs, including adenosine A2A and A2B receptors and dopamine D1 and D5 receptors, did not alter intracellular *Brucella* replication, indicating that exogenous receptor agonism alone is insufficient to recapitulate the infection-associated signaling required for replicative BCV formation. In contrast, genetic or pharmacologic disruption of Gαi-associated signaling enhanced intracellular bacterial growth. Together, these findings indicate that the balance between Gαs- and Gαi-mediated signaling, rather than simple receptor activation, is critical for establishment of the intracellular replicative niche and underscore a central role for cAMP signaling dynamics in BCV maturation.

Although elevated intracellular cAMP is classically linked to activation of protein kinase A (PKA), our data indicate that PKA activity is dispensable for intracellular *Brucella* replication in this system. Pharmacologic inhibition of PKA using a highly specific peptide inhibitor had no effect on intracellular growth, whereas the commonly used small-molecule inhibitor H89 inhibited both intracellular and axenic bacterial growth to a similar extent, consistent with direct antibacterial activity rather than host-directed signaling. These findings argue against a role for PKA as a mediator of the cAMP-dependent effects observed here and instead point to cAMP-dependent, PKA-independent signaling pathways as critical determinants of intracellular permissiveness.

In addition to PKA, cAMP directly activates exchange proteins activated by cAMP (Epac), which signal through Rap family GTPases and regulate vesicular trafficking, cytoskeletal dynamics, autophagy-associated processes, and phagosome maturation (34–36). Moreover, cAMP signaling is highly compartmentalized through localized phosphodiesterase activity, such that inhibition of distinct PDE families can produce divergent biological outcomes despite similar global changes in intracellular cAMP levels (37). In this context, our observation that pharmacologic inhibition of PDE3, and to a lesser extent PDE7, selectively restricts *Brucella* intracellular replication is most consistent with a model in which localized cAMP signaling, rather than bulk cAMP elevation or PKA activation, regulates BCV fate.

Time-course experiments using NanoBRET-based biosensors revealed that Gαs activation occurs at the earliest measurable stages of infection and is sustained over time in a T4SS-dependent manner. Importantly, intracellular replication was sensitive to GPCR pathway disruption only during early infection, whereas inhibition at later time points, after formation of the ER-derived replicative BCV, had little effect. These observations align with established timelines of BCV maturation, in which initial interactions with the ER membrane occur approximately 4-6 h post infection, followed by onset of productive intracellular replication (3, 38, 39). Thus, GPCR-mediated signaling appears to be critical for establishing, rather than maintaining, the replicative BCV.

Consistent with this temporal model, pharmacologic disruption of GPCR-cAMP signaling prior to replicative BCV formation redirected wild-type BCVs into LAMP-1- and p62-positive compartments with features of autophagolysosomal degradation and was accompanied by activation of autophagy-associated signaling pathways. Although whole-cell biochemical analyses preclude definitive assignment of these signaling changes to infected compartments alone, the concordance between altered BCV trafficking, enhanced colocalization with autophagy-associated markers, and failure of intracellular replication supports a model in which inhibition of early GPCR signaling diverts BCVs away from the ER-derived replicative pathway and into degradative compartments.

Consistent with our prior work, sustained phosphorylation of p38 MAP kinase was observed during productive wild-type infection but not during infection with a T4SS-deficient mutant. Notably, p38 phosphorylation was reduced at late time points following pharmacologic disruption of GPCR-cAMP signaling, placing p38 activation downstream of inhibitor-sensitive signaling events and providing temporal context for the signaling cascade described here and previously (6). These observations support a role for p38 activation as a downstream correlate of successful intracellular niche establishment rather than as a primary driver of BCV maturation.

A limitation of this study is that the *Brucella* effector proteins or metabolites responsible for activating host Gαs signaling during infection have not yet been identified. It remains unknown whether *Brucella* directly modifies host G proteins through mechanisms analogous to those employed by classical bacterial toxins such as cholera toxin (40, 41). GPCR-mediated cAMP signaling has also been implicated in macrophage polarization and suppression of antimicrobial effector functions (42), potentially creating a more permissive intracellular environment for replicative BCV establishment, although direct connections between these processes and BCV trafficking remain to be defined.

In summary, our findings support a model in which *Brucella* activates host GPCR-mediated Gαs signaling during early infection, leading to cAMP-dependent signaling events that are independent of protein kinase A (PKA) and are required for maturation of the ER-associated replicative BCV. Pharmacologic disruption of this pathway using ENBA or cilostamide interferes with early signaling dynamics, diverts BCVs into degradative compartments, and results in failure to establish a productive intracellular replicative niche. These results highlight the importance of host GPCR signaling dynamics in *Brucella* pathogenesis and identify Gαs-linked signaling pathways as critical determinants of intracellular survival and potential targets for host-directed therapeutic intervention.

## Materials and Methods

### Bacterial strains, cell lines, and reagents

Bacterial strains used in this study are listed in **Table S4**. *Brucella neotomae* (Bn) and *Legionella pneumophila* (Lp) strains were incubated at 37 °C in a humidified 5% CO₂ incubator on trypticase soy broth (TSB; BD, Franklin Lakes, NJ) supplemented with 50 µg/mL nourseothricin or on buffered charcoal yeast extract (BCYE) medium (43) supplemented with 100 µg/mL thymidine, respectively.

THP-1 human monocyte (ATCC TIB-202, Manassas, VA) and J774A.1 murine macrophage (ATCC TIB-67, Manassas, VA) cell lines were maintained in RPMI 1640 medium (Corning, Tewksbury, MA) supplemented with iron-supplemented calf serum (Gemini Bio-Products).

Pharmacologic inhibitors and agonists used in this study included: DMAB-anabaseine dihydrochloride (Alomone Labs, Jerusalem, Israel); ZM226600, N6-cyclohexyladenosine, SDZ WAG 994, MDL-73005EF hydrochloride, (±)-5′-chloro-5′-deoxy-ENBA, and ABT-724 trihydrochloride (Santa Cruz Biotechnology, Dallas, TX); 8-cyclopentyl-1,3-dimethylxanthine (Fisher Scientific, Waltham, MA); avanafil, cGMP, 8-CPT-cGMP, and UK-432097 (Sigma-Aldrich, St. Louis, MO); cilostazol, cilostamide, and rolipram (Enzo Life Sciences, Farmingdale, NY); H89 dihydrochloride (Selleck Chemicals, Houston, TX); odapipam, A-68930, SKF-38393, BRL-50481, CGS-21680 hydrochloride, and regadenoson hydrochloride (MedChemExpress, Monmouth Junction, NJ); dibutyryl-cGMP, 8-CPT-cAMP, and 8-Br-cAMP (Abcam, Cambridge, MA); 8-Br-cGMP (Tocris Bioscience, Minneapolis, MN); and DPCPX and PKI (14–22) amide, myristolated (trifluoroacetate salt) (Cayman Chemical, Ann Arbor, MI).

### Bacterial intracellular growth assay

J774A.1 murine macrophages or THP-1 human macrophages were seeded at 1 × 10^4^ cells per well in 96-well tissue-culture-treated microplates in a final volume of 100 µL per well and incubated for 24 h at 37 °C with 5% CO₂. Compounds were dispensed using an HP D300 digital dispensing system (TECAN, Morrisville, NC) as previously described (44). Bacterial inocula were prepared in RPMI medium and added to achieve a multiplicity of infection (MOI) of 10 bacteria per host cell.

Plates were centrifuged at 930 × g for 10 min to synchronize infection. At 4 h post infection, gentamicin (100 µg/mL) was added for 1 h to eliminate extracellular bacteria. Wells were then washed twice with PBS and replenished with fresh RPMI medium. Compounds were re-added to maintain the same concentrations present prior to washing. Intracellular bacterial growth was quantified by measuring luminescence at indicated time points using an Infinite M1000 PRO plate reader (TECAN).

### siRNA knockdown experiments

J774A.1 cells were seeded at 5 × 10⁵ cells per well in 6-well plates and incubated overnight. Cells were transfected with 30 nM siRNA (**Table S5**; Integrated DNA Technologies) using Lipofectamine RNAiMAX (Invitrogen) according to the manufacturer’s instructions. Briefly, siRNA-Lipofectamine complexes were prepared in Opti-MEM medium (Thermo Fisher Scientific), incubated for 20 min at room temperature, and then added dropwise to cells cultured in RPMI 1640 supplemented with 10% iron-supplemented calf serum.

At the indicated time points post transfection (6 h or 48 h), cells were harvested and reseeded at 1 × 10⁴ cells per well in 96-well black microplates. Twenty-four hours later, siRNA-transfected cells were infected with Bn, followed by measurement of intracellular cAMP concentrations or intracellular bacterial growth.

### Quantification of intracellular cAMP concentrations

Intracellular cAMP levels were quantified using the cAMP-Glo assay (Promega, Madison, WI) according to the manufacturer’s instructions. At indicated time points, cells were washed twice with PBS and lysed with 20 µL cAMP-Glo lysis buffer, followed by incubation for 20 min at room temperature. Forty microliters of cAMP detection solution containing protein kinase A was added, followed by 60 µL of Kinase-Glo reagent. Luminescence was measured using an Infinite M1000 PRO plate reader.

### NanoBRET analysis

NanoBRET vectors were constructed using primers listed in **Table S4**. Full-length eYFP (717 bp), mouse Gαs (1,182 bp), and Gγ (204 bp) sequences were synthesized as gBlocks (Integrated DNA Technologies). Gγ was fused to the C-terminus of eYFP via overlap PCR, with an upstream IRES to permit eukaryotic expression, and cloned into the pRetroX vector using BamHI and EcoRI restriction sites, yielding pRetroX-IRES-eYFP::Gγ. NanoLuc luciferase was fused to the C-terminus of Gαs with an upstream IRES and cloned into the same vector using NotI and BamHI, yielding pRetroX-Gαs::NLuc-IRES-eYFP::Gγ.

Plasmids were purified using a Qiagen miniprep kit and transfected into J774A.1 cells at 60-70% confluency using Lipofectamine LTX with PLUS reagent (Invitrogen), with 1 µg DNA per well. After 48 h, transfected cells were washed with PBS and seeded at 1 × 10^4^ cells per well in 96-well white plates and allowed to adhere overnight. Bacteria were added at an MOI of 50, followed by centrifugation at 930 × g for 10 min.

At indicated time points, NanoLuc substrate was added, and NanoBRET signals were measured using the Infinite M1000 PRO plate reader. BRET was detected between NanoLuc (460/12 nm) and eYFP (530/12 nm), with an integration time of 0.5 s. BRET ratios were calculated as the ratio of acceptor (530 nm) to donor (460 nm) emission.

### Western blot analysis

J774A.1 macrophages were seeded at 5 × 10^5^ cells per well in 6-well plates and infected at the indicated MOI. For short infections (0.5 h and 2 h), gentamicin was not added. For 24 h and 48 h infections, gentamicin was added at 4 h post infection as described above. Cells were lysed in RIPA buffer supplemented with protease and phosphatase inhibitors (Cell Signaling Technology), clarified by centrifugation at 12,000 × g for 10 min at 4 °C, and protein concentrations were determined by BCA assay.

Samples (5-10 µg protein) were resolved on 8-16% SDS-PAGE gradient gels, transferred to nitrocellulose membranes, blocked in TBST with 5% BSA, and probed with primary antibodies (1:1,000) overnight at 4 °C. Secondary antibodies (1:10,000) were applied for 1.5 h at room temperature. Detection was performed using SuperSignal West Femto substrate and imaged on a Bio-Rad ChemiDoc system.

### Confocal imaging

Confocal microscopy was performed as previously described (16). J774A.1 cells were seeded at 5 × 10^4^ cells per well on 12-mm coverslips in 12-well plates. Cells were treated with inhibitors and infected with tdTomato-expressing Bn at an MOI of 10. After synchronized infection and gentamicin treatment, cells were fixed, permeabilized, blocked, and stained with primary antibodies (1:100) and Alexa Fluor-conjugated secondary antibodies (1:300). Images were acquired on a Zeiss LSM 880 confocal microscope using Zen 2.1 software for colocalization analysis.

### Statistical analysis

Statistical analyses were performed using GraphPad Prism 10. Data represent two or three independent biological replicates. IC₅₀ values were calculated using three-parameter nonlinear regression. Statistical significance was determined by one-way ANOVA with Dunnett’s post hoc test. *P* < 0.05 was considered significant.

## Funding and Acknowledgements

This work was supported by the National Institute of Allergy and Infectious Diseases of the National Institutes of Health under award number R01AI099122 to J.E.K. and by a Novel Therapeutics Development Grant from the Massachusetts Life Sciences Center. The content is solely the responsibility of the authors and does not necessarily represent the official views of the National Institutes of Health. We thank TECAN for use of the Infinite Pro M1000 and D300 instrumentation. TECAN had no role in study design, manuscript preparation, or decision to publish.

