## Supplementary Tables and Figures for "Host GPCR-cAMP signaling balances Gαs and Gαi activity to control intracellular *Brucella* infection"

Table S1. The effects of Adenosine or dopamine receptor agonists on Bn intracellular growth in THP-1 macrophages.[a]

| Target & Functions | Compounds | Z-score |
|---|---|---|
| Adenosine A1 receptor agonist | N6-Cyclopentyladenosine; CPA | -7.0 (Strong) |
| | 2-Chloro-N6-cyclopentyladenosine | -4.3 (Weak) |
| | N6-Cyclohexyladenosine; CHA | -7.1 (Strong) |
| | SDZ WAG 994 | -7.0 (Strong) |
| | (±)-5'-Chloro-5'-deoxy-ENBA | -7.2 (Strong) |
| | 2'-MeCCPA | -1.2 |
| Adenosine A3 receptor agonist | 2-Cl-IB-MECA | -3.1 (Weak) |
| | IB-MECA | -2.5 |
| | Inosine | -1.9 |
| Adenosine A2A receptor agonist | CGS-21680 | -0.3 |
| | Limonene | -2.0 |
| Adenosine A2B receptor agonist | NECA | 1.1 |
| Dopamine D1 & D5 receptor agonist | Dihydrexidine | -3.0 |
| | Dopamine | 1.9 |
| | Fenoldopam | -0.89 |
| Dopamine D1 receptor agonist | SKF-82958 | 2.7 |
| | SKF-38393 | 6.5 |
| | 6-Br-APB | -0.17 |

| | | |
|---|---|---|
| | A-68930 | 5.6 |
| | Cabergolin | 1.9 |
| | Pergolide | 2.0 |
| Dopamine D4 receptor agonist | WAY-100635 | -6.3 (Medium) |
| | ABT 724 trihydrochloride | -7.3(Strong) |
| | PD 168077 maleate | -6.9 (Medium) |
| | CP-226269 | -1.1 |
| Dopamine D4 receptor antagonist | Clozapine[b] | 10.0 |
| Serotonin 5-HT1A receptor agonist | MDL 73005EF hydrochloride | -9.5 (Strong) |

[a]Data from Kang & Kirby (1). Z-scores were calculated across screening plates from the primary THP-1 macrophage screen to identify statistically significant inhibitors of intracellular *B. neotomae* growth.

[b]Mixed agonist/antagonist. Strongest effects are as D4 antagonist (2).

Table S2. Phosphodiesterase (PDE) inhibitors affecting Bn intracellular growth in THP-1 macrophages.[a]

| Compounds | Primary PDE Inhibitory Activity (reported) | Bn inhibition (% of control) |
|---|---|---|
| IBMX | nonselective | -4.8% |
| Pentoxifyline | nonselective (weak) | -9.8% |
| Papaverine | nonselective | -27% |
| Ibudilast | nonselective | -21% |
| Vinpocetine | PDE1 (additional targets described) | 28% |
| Cilostazol | PDE3 | 90% |
| Cilostamide | PDE3 | 89% |
| Enoximone | PDE3 | 83% |
| Imazodan | PDE3 | 40% |
| Milrinone | PDE3 | 2.4% |
| Trequinsin | PDE3 | 45% |
| Olprinone | PDE3 | 33% |
| Quazinone | PDE3 | 12% |
| ICI-63197 | PDE3/4 | 20% |
| YM 976 | PDE4 | 24% |
| Rolipram | PDE4 | -0.7% |
| Quercetin | PDE4 (pleiotropic flavonoid) | -64% |
| Sildenafil | PDE5 | -3.5% |
| Zaprinast | PDE5 | -0.5% |
| BRL 50481 | PDE7 | 53% |

[a]Data from Kang & Kirby (1). Percent inhibition values reflect normalized intracellular growth relative to vehicle-treated controls and are provided to convey biological effect size.

**Table S3. Axenic growth $IC_{50}$ values, host cell cytotoxicity, and maximal rescue of ENBA-mediated inhibition of intracellular *Brucella neotomae* growth by cyclic nucleotide analogs.**

| | 8-Br-cAMP | 8-CPT-cAMP | 8-Br-cGMP | Dibutyryl-cGMP | cGMP | 8-CPT-cGMP |
|---|---|---|---|---|---|---|
| Bn axenic growth $IC_{50}$ (µg/mL) | >50 | >50 | >50 | >50 | >50 | >50 |
| Host cell $CC_{50}$ (J774A.1, µg/mL) | 25 | 32 | >50 | 16 | >50 | >50 |
| Maximal fold rescue of intracellular growth | 15 | 10 | No rescue | 2.3 | 2.0 | No rescue |
| Concentration alleviating ENBA-mediated inhibition (µg/mL) | 21 | 21 | No rescue | 9.1 | 50 | No rescue |

Data shown correspond to rescue experiments presented in Fig. 3C. Cyclic nucleotide analogs were tested in J774A.1 macrophages infected with *B. neotomae* in the presence of ENBA (2 µg/mL). Maximal fold rescue reflects the increase in intracellular luminescence relative to ENBA-treated controls. Concentrations listed were below axenic growth $IC_{50}$ values and host cell cytotoxicity thresholds.

Table S4. Bacterial strains, cell lines, and primers used in this study.

| ***Bacterial strains*** | | |
|---|---|---|
| *Strain* | *Relevant characteristics* | *Source or Reference* |
| *B. neotomae* 5K33 | Parent biosafety level 2 rodent pathogen | BEI Resources |
| *B. neotomae*-Lux | Transposon mutant of 5K33 expressing Lux operon | (3) |
| *B. neotomae*-tdTomato | Transposon mutant of 5K33 having proD/tdtomato-nat genes | |
| *B. neotomae* ΔvirB4-Lux | Transposon mutant of virB4 in-frame deletion mutant expression Lux-operon | |
| *B. neotomae* ΔvirB4-tdTomato | Transposon mutant of Bn ΔvirB4 having proD/tdtomato-nat genes | |
| *L. pneumophila* 02fla-Lux | *L. pneumophila* 02fla having Lux operon | (4) |
| NEB-5α | *fhuA2 Δ(argF-lacZ)U169 phoA glnV44 Φ80 Δ(lacZ)M15 gyrA96 recA1 relA1 endA1 thi-1 hsdR17* | NEB |
| ***Eukaryotes*** | | |
| *Cell line* | | *Source or Reference* |
| J774A.1 | | ATCC TIB-67 |
| THP-1 | | ATCC TIB-202 |
| ***Oligonucleotides*** | | |
| *Name* | *Sequences* | *Characteristics* |
| IRES-F | CGC*GGATCC*CCCCTCTCCCTCCCCC | IRES amplification & IRES-eYFP fusion |
| IRES_eYFP-R | CATCATGGTGGCTTATCATCGTGTTTTCA | |
| eYFP-F | ATAAGCCACCATGATGGTGAGCAAGGGCGA | eYFP amplification & IRES-eYFP::Gγ fusion |
| eYFP::Gγ-R | CGC*GAATTC*CTAGAGAATTATGCAAGGCTT | |
| Gαs-F | CGC*GCGGCCGC*GCCACCATGGGCTGCCTCGG | Gαs subunit amplification & Gαs::NLuc fusion |
| Gαs::NLuc-R | GGAGCCGCCACCACCGAGCAGCTCGTATTG | |
| NLuc-F | CTCGGTGGTGGCGGCTCCGTCTTCACACTCGAA | NLuc amplification & Gαs::NLuc fusion |
| NLuc-R | CGC*GGATCC*TTACGCCAGAATGCGTTCGCA | |

Table S5. siRNA used in this study.

| Target | Sequences | Use |
|---|---|---|
| Adenosine A1 receptor | Sense 5'-AGCAUGGAGUACAUGGUCUACUUCA-3' | Adenosine A1 knockout |
| | Antisense | |

| | | |
|---|---|---|
| | 5’-UGAAGUAGACCAUGUACUCCAUGCUGA-3’ | |
| Dopamine D4 receptor | Sense<br>5’-GCAGACACCCACCAACUACUUCATC-3’ | Dopamine D4 knockout |
| | Antisense<br>5’-GAUGAAGUAGUUGGUGGGUGUCUGCAG-3’ | |
| Non-target siRNA | Sense<br>5′-UUCUCCGAACGUGUCACGU-3′ | |
| | Antisense<br>5′-ACGUGACACGUUCGGAGAA-3′ | |

### Supplementary Figures

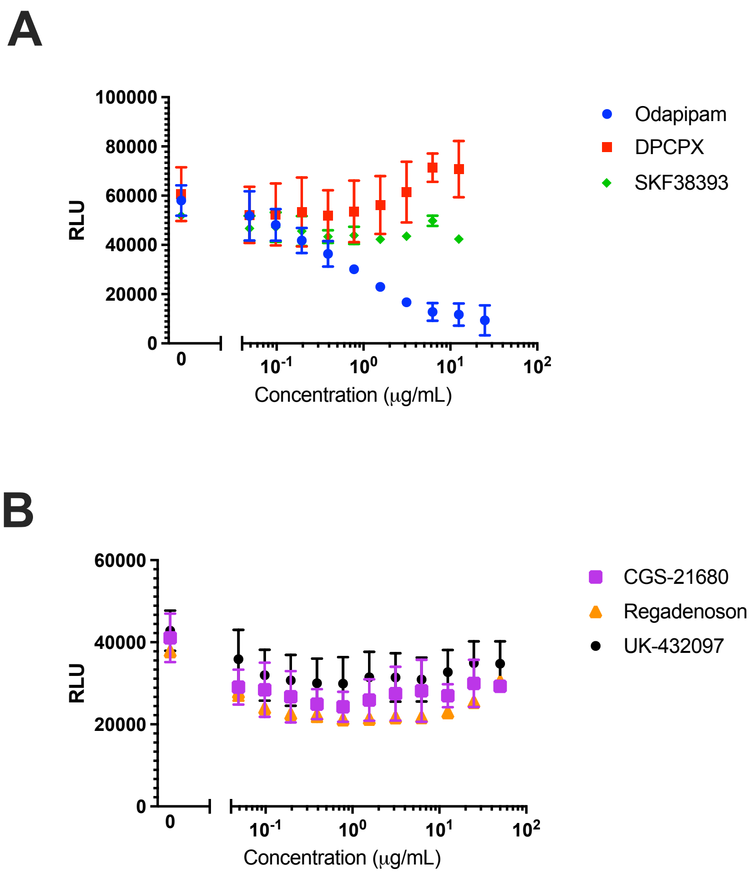

**Figure S1. GPCR pathway specificity of host-mediated modulation of *B. neotomae* intracellular growth. (A)** Intracellular growth of luminescent *B. neotomae* (Bn) in J774A.1 macrophages measured 48 h post infection following treatment with ligands targeting dopamine and adenosine receptors with distinct G-protein coupling profiles. Odapipam (dopamine D1/D5 receptor antagonist; Gαs-associated), SKF38393 (dopamine D1/D5 receptor agonist; Gαs-associated), and DPCPX (adenosine A1 receptor antagonist; Gαi-associated) were tested over a range of concentrations. **(B)** Intracellular Bn growth measured 48 h post infection following treatment with selective adenosine A2A receptor agonists CGS-21680, regadenoson, and UK-432097, all of which signal predominantly through Gαs. Across the concentrations tested, including the highest concentrations, activation of Gαs-coupled adenosine A2A receptors or dopamine D1/D5 receptors did not significantly alter intracellular Bn growth, whereas antagonism of the Gαs-coupled dopamine D1/D5 receptor with odapipam resulted in dose-dependent inhibition. Data points represent the mean ± standard deviation (SD) calculated from single measurements obtained in two independent experiments. Intracellular bacterial burden is reported as relative luminescence units (RLU).

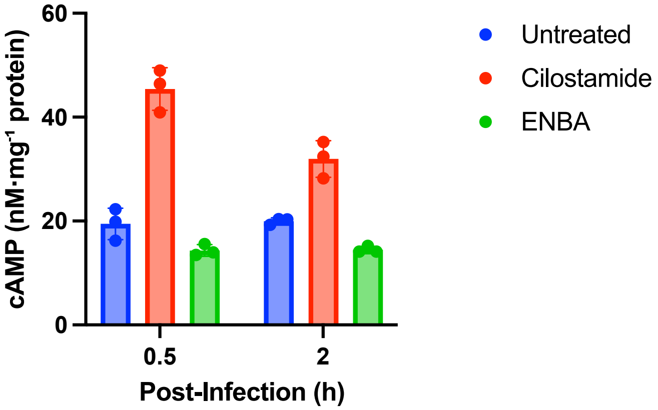

**Figure S2. Effects of cilostamide and ENBA on intracellular cAMP levels in uninfected J774A.1 macrophages.** Uninfected J774A.1 macrophages were treated with the PDE3 inhibitor cilostamide or the adenosine $A_1$ receptor agonist ENBA (each at 2 µg/mL), and intracellular cAMP levels were measured after 30 min and 2 h. cAMP concentrations were normalized to total cellular protein. Data represent the mean ± SEM from three independent experiments.

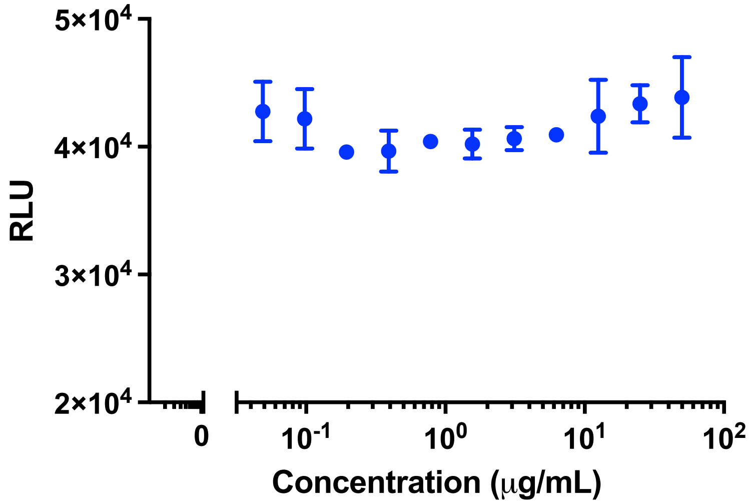

**Figure S3.** The specific PKA inhibitor PKI 14-22 amide does not affect intracellular *Brucella neotomae* growth. J774A.1 macrophages were infected with luminescent *B. neotomae* and treated with increasing concentrations of the myristoylated PKA inhibitor PKI 14-22 amide. Intracellular bacterial growth was quantified by luminescence 48 h post-infection. No inhibition of intracellular growth was observed, including at the highest concentration tested (50 μg/mL). Data represent the mean ± standard deviation of two independent measurements.

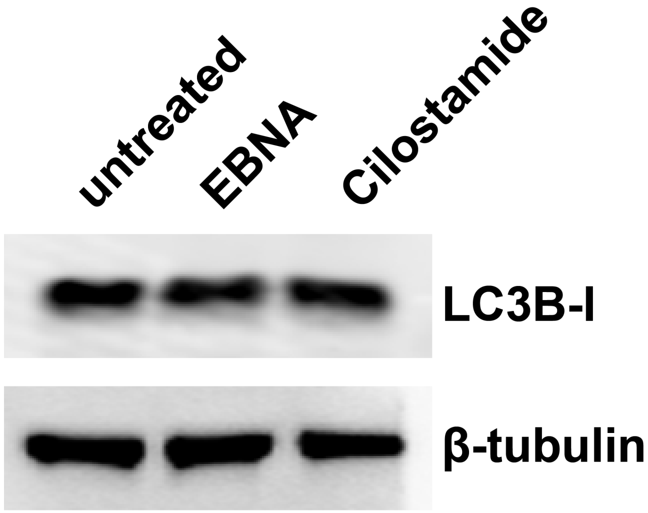

**Figure S4. Effects of cilostamide and ENBA on LC3B levels during ΔvirB4 infection of J774A.1 macrophages.** J774A.1 macrophages were infected with the type IV secretion system-deficient ΔvirB4 mutant of *Brucella neotomae* and treated with cilostamide or ENBA (2 µg/mL). After 48 h, LC3B levels were assessed by immunoblotting. Only the non-lipidated form, LC3B-I, was detected under these conditions; the lipidated form LC3B-II was not observed. β-tubulin served as a loading control. These findings indicate that modulation of intracellular replication by ENBA or cilostamide is not associated with detectable LC3B lipidation under ΔvirB4 infection conditions.

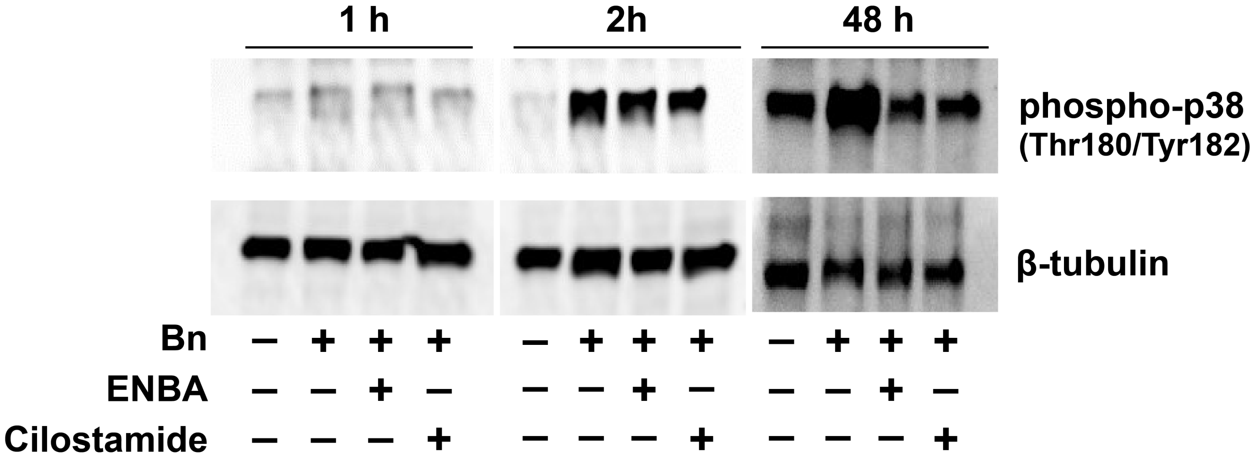

**Figure S5. Effects of ENBA and cilostamide on p38 MAP kinase phosphorylation during *Brucella neotomae* infection.** J774A.1 macrophages were infected with wild-type *B. neotomae* and treated with ENBA or cilostamide (2 µg/mL), as indicated. Cell lysates were collected at early (1 h and 2 h) and late (48 h) time points post infection, and phosphorylation of p38 MAP kinase was assessed by immunoblotting. β-tubulin served as a loading control. During early infection, p38 phosphorylation was induced by *B. neotomae* infection and was not altered by ENBA or cilostamide treatment. In contrast, at 48 h post infection, p38 phosphorylation was reduced by both ENBA and cilostamide.
